# HMCVelo: A Deterministic Model for Hydroxymethylation Velocity in Single Cells

**DOI:** 10.64898/2026.04.20.719607

**Authors:** Paramita Mishra

## Abstract

I present *hydroxymethylation velocity* (HMCVelo), the first velocity framework for DNA methylation dynamics. HMCVelo is a deterministic ordinary differential equation (ODE) model that computes the time derivative of hydroxymethylation state for individual cells and genes. The model exploits a recent advance in single-cell epigenomics—Joint single-nucleus hydroxymethylcytosine and methylcytosine sequencing (Joint-snhmC-seq)—which enables subtraction-free quantification of 5-hydroxymethylcytosine (5hmC) and 5-methylcytosine (5mC) at single-cell resolution, resolving temporal methylation dynamics from static molecular snapshots. HMCVelo models the methylation–demethylation cycle as three coupled processes—methylation, hydroxymethylation, and demethylation—governed by gene-specific rate parameters estimated at steady state via constrained least-squares regression. Scale invariance reduces the parameter space from three to two free parameters per gene. Applied to murine cortical cells (*n* = 519 and *n* = 545), HMCVelo infers cellular trajectories with velocity confidence scores exceeding 0.89 across all cell types, compared to confidence scores below 0.45 when RNA velocity is repurposed on the same data. I further prove that in any closed biochemical system with a conservation law, the complement variable cannot resolve trajectory bifurcations—a result with implications for embedding basis selection in all future velocity frameworks applied to cyclic biochemical systems. This work provides a foundation for multi-omic trajectory inference integrating epigenetic and transcriptomic measurements.

## 1 Introduction

Every cell carries, in the chemical modifications of its DNA, a continuous record of where it has been and a set of instructions for where it is going. To read this record directly — rather than through the downstream proxies of gene expression — is to observe differentiation at its point of origin.

Cell differentiation in neural cells was traditionally viewed as a transition between discrete, static cell types. In actuality, there exist continuous spectra of cell states in differentiating cells, characterized by two aspects: a direction toward future states and a current position relative to the initial state in a differentiation path. Trajectory inference (TI) methods assign cells a pseudotime value along these paths, capturing both directionality and relative position within a lineage [1]. Since pseudotime is based solely on molecular information, it may not directly equal chronological time. However, it is assumed that the ratio of two cellular pseudotimes correlates with the ratio of their respective chronological times since the beginning of differentiation. The trajectory is then defined as a collection of cellular lineages, together defining the dynamic differentiation process.

5-hydroxymethylcytosine (5hmC), an oxidized form of 5-methylcytosine (5mC) generated by the enzymatic activity of ten-eleven translocation (TET) family of DNA dioxygenases, displays dynamic cell- and gene-specific heterogeneity, potentiating its functional role in cellular trajectory specification [2, 3]. While the exact function of 5hmC is still being determined, there are several reasons why 5hmC dynamics must be investigated in differentiating brain cells. 5hmC is the most abundant oxidized form of 5mC and shows stable enrichment at certain genomic sites [4]. In the brain specifically, levels of global 5hmC increase from birth to adulthood [5], and 5hmC crosstalk with histone marks may play a causal role in neuronal maturation and age-related neurodegenerative processes such as Alzheimer’s disease [6]. Preferential 5hmC enrichment in the gene bodies of expressed genes and 5hmC abundance in exons is positively correlated with RNA expression in the murine brain in a cell type–specific manner [7]. Empirically, 5hmC events correlate in time and space with developmental stages in differentiating cells [4].

Bulk DNA methylome profiling analyses have hinted at the heterogeneity of 5hmC in brain cells but cannot resolve epigenetic heterogeneity and molecular dynamics correlated with distinct cell states [8]. Previous methods to measure methylation dynamics have been limited to non-cell-specific sequencing techniques or indirect inference using transcriptomic and chromatin data. Despite improvements in single-cell epigenetics protocols, 5hmC and 5mC rates have historically been unresolved at single-cell level. Single-cell bisulfite sequencing (scBS-seq) has revolutionized the study of single-cell DNA methylomes but cannot resolve the base ambiguity between 5hmC and 5mC [9]. Probabilistic models using 5hmC enrichment have inferred lineage trees at cell-division resolution but cannot determine directionality or rates of methylation state transitions [10].

The recent protocol ACE-seq uses DNA deaminase APOBEC3A (A3A) for deamination of unmodified C, 5fC, 5caC, and 5mC, but preserves 5hmC due to enzymatic protection by glucosylation [11]. Based on this approach, Wu and colleagues developed Joint-snhmC-seq, which combines bisulfite treatment with A3A deamination to generate subtraction-free counts of 5hmC using bisulfite-treated ACE sequencing (Fig. 1). In parallel, the total level of DNA methylation (5mC+5hmC) is profiled using single-cell BS-seq [12]. This joint epigenetic profiling strategy, coupled with the consistency of methylation–demethylation mechanisms, provides an opportunity to theorize a vector representing methylation dynamics using a deterministic model.

**Figure 1:**
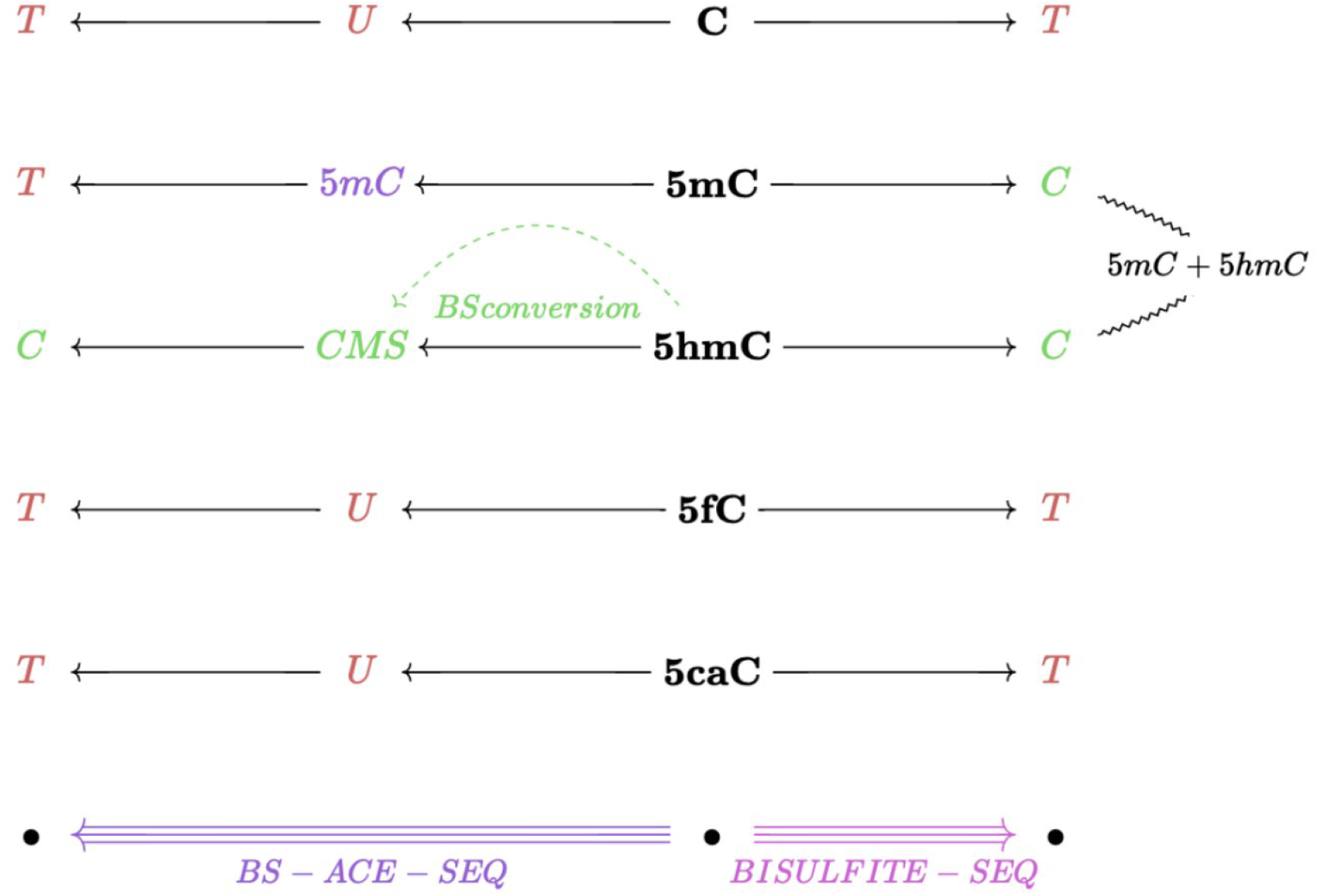
Conversion of unmodified and modified cytosines in scBS-seq (pink) and BS-treated ACE-seq (purple). scBS-ACE-seq operates through the conversion of 5hmC to cytosine-5-methylenesulfonate (CMS). Rare oxidized forms of 5mC (5fC and 5caC) are deaminated in the process, resulting in subtraction-free counts of 5hmC. Both techniques are used in JointSeq on cell-specific samples.

RNA velocity [13]—the time derivative of gene expression state—is a deterministic model which exploits the difference in spliced and unspliced single-cell mRNA counts to capture expression dynamics, producing a high-dimensional vector which can be leveraged to predict directions and transition probabilities of cellular transitions. Several related models have since been developed: a dynamical version of RNA velocity [14], chromatin velocity for chromatin dynamics [15], protein velocity extending steady-state RNA velocity to protein measurements [16], and multi-omic velocity integrating chromatin and gene expression dynamics [17]. However, at present, there are no such models for DNA methylation dynamics.

In its simplest form, RNA velocity can potentially be repurposed for temporal DNA methylation dynamics by substituting unspliced mRNA for 5mC and spliced mRNA for 5hmC. However, while RNA splicing is a unidirectional process, methylation occurs cyclically due to the action of TET and DNMT enzymes on CpG dinucleotides (Fig. 2). Furthermore, unlike mRNA, there is no explicit degradation of cytosine, calling for the inclusion of unmodified cytosine levels to model velocity vectors more precisely. A model more specific to methylation and demethylation dynamics may allow for a better mechanistic understanding of cellular trajectories, and reveal the interplay of epigenetics and gene expression in modulating cell state.

**Figure 2:**
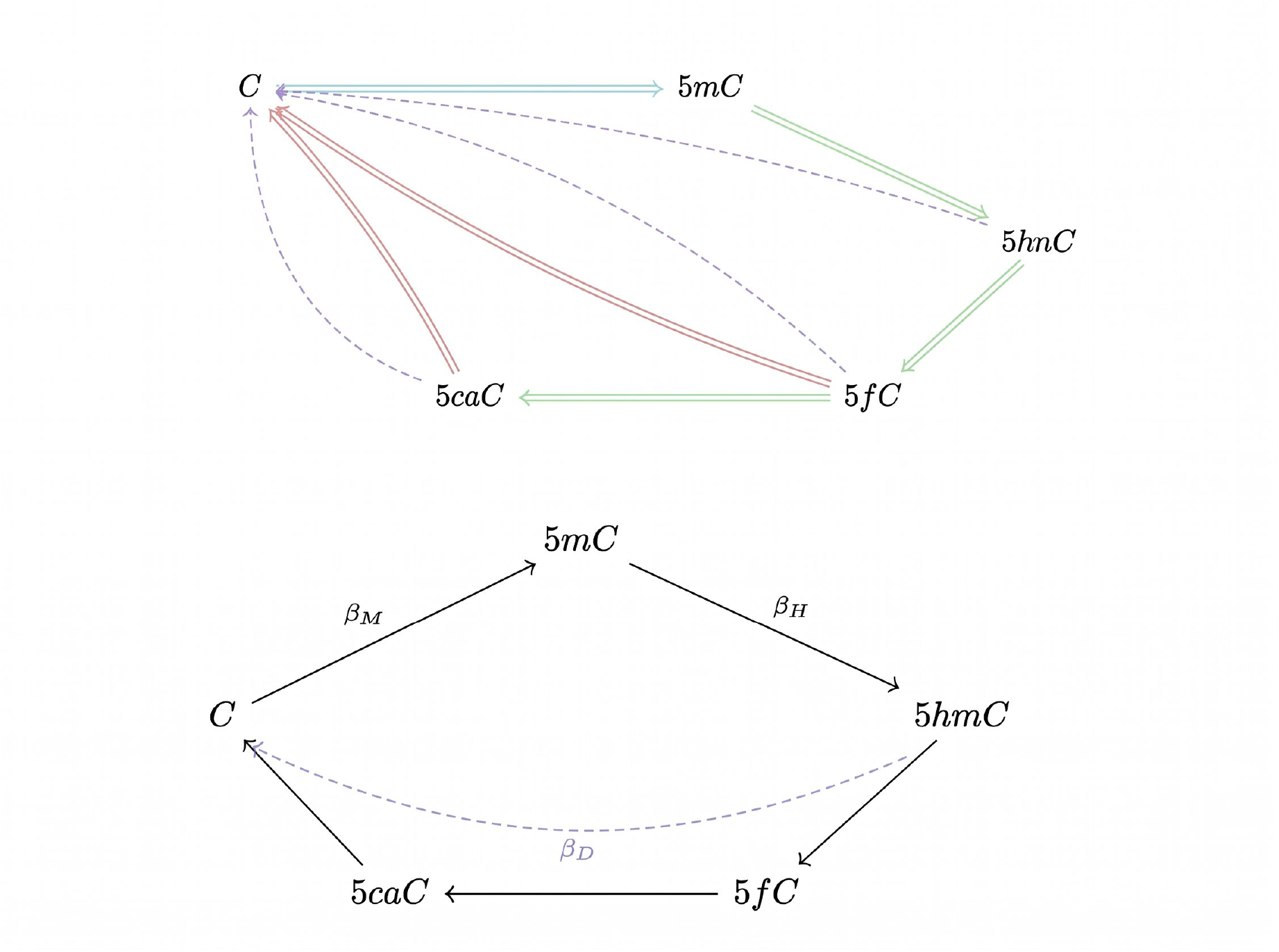
(a) Demethylation cycle: Active DNA demethylation can occur through active modification–passive dilution (AM–PD) or active modification–active removal (AM–AR). TET enzyme conversion is shown in green, DNMT in blue. (b) Demethylation cycle modeled by HMCVelo rate parameters.

Here I introduce HMCVelo, the first velocity model for DNA methylation dynamics. HMCVelo infers the *hydroxymethylation velocity* —the time derivative of hydroxymethylated cytosine—of cells. To quantify the 5mC and 5hmC dynamics, the methylation–demethylation cycle is categorized into three distinct processes: methylation, hydroxymethylation, and de-methylation (Fig. 2b). These processes are modeled using ordinary differential equations and solved numerically. I demonstrate that HMCVelo resolves lineage branching and cellular trajectories in murine cortical neurons with substantially higher confidence than RNA velocity applied to the same methylation data, and identify candidate differentiation-associated genes ranked by velocity dynamics.

## 2 Model

### 2.1 Previous approach: RNA Velocity

The initial RNA velocity model exploited the difference between features of observed unspliced and spliced mRNA reads from scRNA-seq. Labeling pre-mRNA as unspliced (*u*) and post-mRNA as spliced (*s*), the following ODEs are created for each gene and each cell:

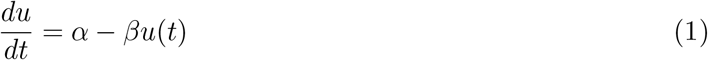

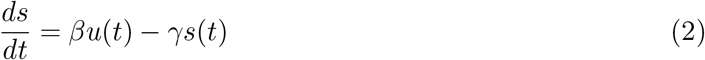

where *γ* is the spliced RNA degradation rate, *β* is the unspliced RNA splicing rate, and *α* is the unspliced RNA transcription rate. RNA velocity is then defined as 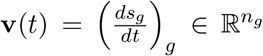. To capture transcriptional switching, *α* is modeled with a switch time *t*_*s*_: *α*(*t*) = *α*^on^ for *t* ≤ *t*_*s*_ and *α*(*t*) = 0 for *t > t*_*s*_. The ODEs can be solved for *s*(*t*) analytically, and the rate parameters can be estimated assuming the system has reached steady state [13, 14].

### 2.2 Model definition

In accordance with the methylation–demethylation cycle (Fig. 2a), HMCVelo is defined by the following three equations, where *c*(*t*) represents the expected quantity of unmodified cytosine, *m*(*t*) represents the expected quantity of methylated cytosine, *h*(*t*) represents the expected quantity of hydroxymethylated cytosine, and rate parameters as described below:

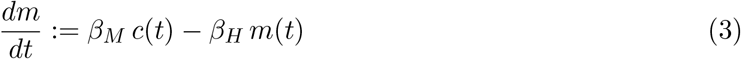

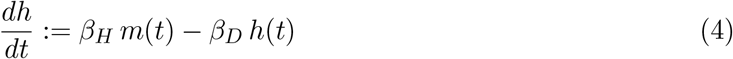

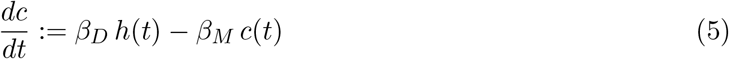

with the following rate parameters representing methylation, hydroxymethylation, and de-[hydroxy]methylation (Fig. 2b) respectively:

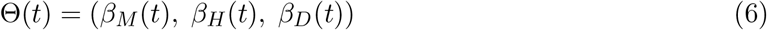

Assuming gene-specific, time-independent rate constants given a short time-step:

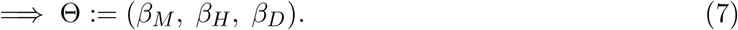

For experiment 1, the observed values are quantities of *h*(*t*), *m*(*t*), and *c*(*t*). For experiment 2, the observed values are the proportions of 5hmC and 5mC, *x*_*m*_(*t*) and *x*_*h*_(*t*) respectively:

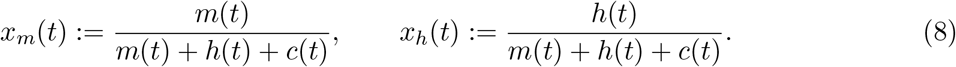

The proportion of cytosine is then inferred as:

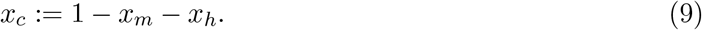

Note that Eq. (5) is linearly dependent on Eqs. (3) and (4), since 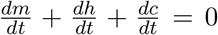, which implies that *m*(*t*) + *h*(*t*) + *c*(*t*) is constant. This conservation law permits equivalent analysis using either absolute quantities or proportions. Since *m*(*t*) + *h*(*t*) + *c*(*t*) is constant, one can substitute *m*(*t*) for *x*_*m*_(*t*), *h*(*t*) for *x*_*h*_(*t*), and *c*(*t*) for *x*_*c*_(*t*) in the original model definition (by dividing both sides by *m*(*t*) + *h*(*t*) + *c*(*t*)), and all the same dynamics follow. Hence, I continue this analysis using *c*(*t*), *h*(*t*), *m*(*t*) without loss of generality.

**Hydroxymethylation velocity, v**(*t*), is the time-derivative of hydroxymethylation with individual components for each gene *g*, such that

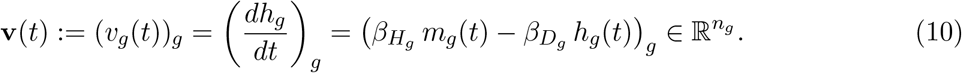

### 2.3 Direction of hydroxymethylation

As compared to methylation, hydroxymethylation may be a better estimate of the direction of transition to a more mature cell state. During neurogenesis, 5hmC increases as cells mature over time [6]. Compared to hydroxymethylation, at a whole-genome (i.e. dimensionality-reduced) level, methylation is more constant, and unmodified cytosine proportion reflects the changes in either of these two methylated cytosines. Therefore, hydroxymethylation is most likely to have a direction parallel to a cellular trajectory.

#### Usage of hydroxymethylation velocity in other tissues

In the brain, hydroxymethylation is a stable and relevant epigenetic mark. For other tissues, demethylation as a whole is more relevant than hydroxymethylation alone. Therefore, when considering this model for other use-cases, it is useful to consider *demethylation velocity* **d**(*t*):

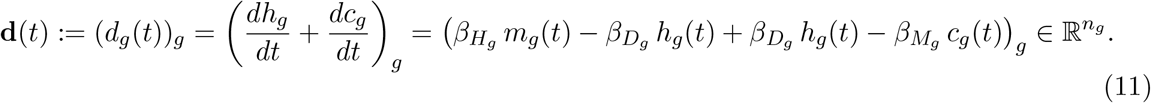

### 2.4 Exclusion of 5fC and 5caC

There are three intermediate states between mC and C during demethylation: 5hmC, 5fC, and 5caC (Fig. 2a). The sequencing techniques bisulfite sequencing and bisulfite-treated ACE-seq cannot resolve 5fC and 5caC. It is safe to exclude 5fC and 5caC in the model due to the following reasons. First, 5fC and 5caC are considered more transient intermediates at any given pseudotime [19], present at levels orders of magnitude lower than 5hmC and 5mC in wild-type cells, as determined by very sensitive methods [20, 21, 22]. Second, the proportions for 5fC and 5caC are implicitly part of the *c*(*t*) variable due to the nature of the sequencing protocol (Fig. 1). *β*_*D*_ is assumed to include all stages of active de-hydroxymethylation to C, assuming some 5fC and 5caC dynamics are recovered by this rate parameter.

### 2.5 Non-active methylation

Methylation dynamics can also occur passively. This process depends on DNA replication [23], and a single-cell snapshot of methylation abundance does not provide adequate information to model passive demethylation. This is assumed to be out of scope for the present model.

### 2.6 Scale invariance

The system exhibits scale invariance. We assume the following for any scaling parameter *κ >* 0:

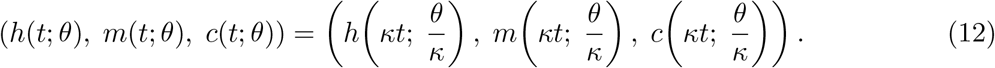

For example, doubling all the rate parameters would result in *h*(*t*), *m*(*t*), *c*(*t*) reaching a particular state in half the time. Under this assumption, we can set one of the rate parameters to a constant value to fix the time scale of the system, noting that the constant we choose will affect the relative magnitude (importantly, not the direction) of HMC velocity **v**. This approach has been used in previous single-cell models for gene expression [24]. For our model, we set *β*_*H*_ = 1, such that:

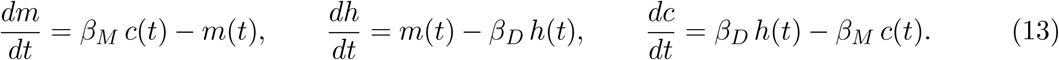

## 3 Parameter Estimation

Rate parameters are estimated on a gene-specific basis at steady state as defined below.

### 3.1 Least-squares regression

In the steady state, when we have:

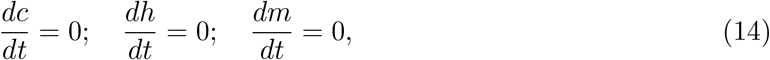

the following equations apply:

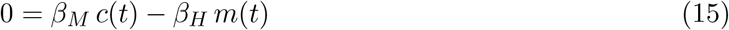

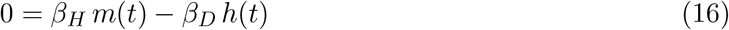

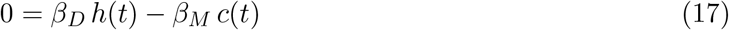

Since Eq. (17) is implied by Eqs. (15) and (16), we only need Eqs. (15) and (16) to estimate rate parameters.

#### Experiment 1

Since experiment 1 uses read counts, single-snapshot observations *c, m, h* are used. Additionally, 100-Kb bin *b* is used for experiment 1 instead of gene *g*: *g* ≡ *b*.

#### Experiment 2

Experiment 2 uses proportions. Recalling from above that the dynamics of *x*_*m*_, *x*_*h*_, *x*_*c*_ are the same as *m, h, c* since *x*_*m*_ + *x*_*h*_ + *x*_*c*_ is constant, the observed values are 1 − *x*_*h*_ − *x*_*m*_, *x*_*m*_, *x*_*h*_.

#### Null value estimates

The rate parameters for any genes with many nulls past the drop stage were returned as [0, 0, 0].

### 3.2 Applying scale invariance

It is noteworthy that due to no offset, one steady-state solution will always be *θ* = (0, 0, 0). However, by the scale invariance property, *β*_*H*_ = 1, so (0, 0, 0) is naturally excluded from our solution space. The `HMCVelo.least_squares_params` function solves for the following hypothesis function.

Setting *β*_*H*_ = 1:

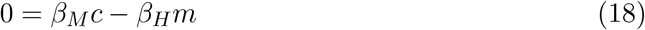

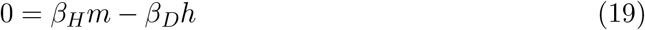

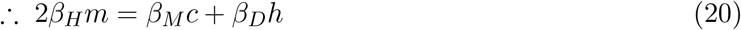

Setting *β*_*H*_ = 1:

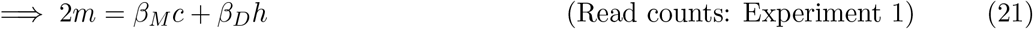

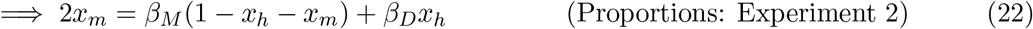

This linear equation is solved via constrained least-squares regression (`scipy.optimize.least_squares` [25]) with no intercept term, using the cost function min 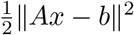 subject to lb ≤ *x* ≤ ub.

#### Gene-specificity of coefficient

*S*_*g*_ is the input to `least_squares_params` for gene *g* and cell *c* such that

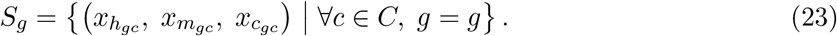

Therefore, coefficients *θ* are gene-specific and cell-independent.

## 4 Computational Implementation

Due to the complexity of the closed-form solution for the model, a numerical rather than analytical solution is implemented, a significant deviation from RNA velocity. The `HMCVelo.solve_model` function resolves ODEs for each gene and each cell by implementing `odeint` from SciPy with initial conditions (*m*_0_, *h*_0_, *c*_0_) set as the observed quantities. The model definition passed to `solve_model` is:

~~~
def model_equations(y, t, betaM, betaH, betaD):
    m, h, c = y
    dm_dt = betaM * c - betaH * m
    dh_dt = betaH * m - betaD * h
    dc_dt = betaD * h - betaM * c
    return [dm_dt, dh_dt, dc_dt]
~~~

The solution of 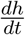 for a time-step of *t* is assumed to be the solved abundance of 5hmC at *t* minus the initial state, divided by *t*. The time-step was set as Δ*t* = 10^−5^ after finding that Δ*t <* 10^−5^ does not significantly change the solution. This is extracted and stored in the solutions array with dimensions *n*_cells_ × *n*_features_.

### 4.1 Dimensionality reduction

HMC Velocity is a high-dimensional vector requiring dimensionality reduction (DR) for qualitative interpretation. Using scVelo [14], incremental PCA and non-linear reduction using t-SNE and UMAP were implemented. DR was implemented on *observed methylation states* (OMS, passed as `AnnData.X`) which represents the sum of 5hmC and 5mC for each cell and each feature. This choice was inspired by RNA Velocity, which uses total observed expression state. OMS is equivalent to the subtraction-free scBS-seq data (Fig. 1).

### 4.2 Velocity embeddings

When cell states are embedded in a low-dimensional space using DR, one could assume the extrapolated cell state using HMCVelo could be used to plot a vector arrow to the “next” state for linear reduction. For non-linear t-SNE and UMAP, this assumption is unsubstantiated. To create a robust velocity embedding, scVelo’s velocity embedding method is implemented.

Assuming difference in embedding positions is calculated as 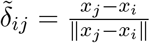, and transition probabilities aggregated into a transition matrix 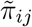 describing the Markov chain for the differentiation, expected displacements with respect to the transition matrix are:

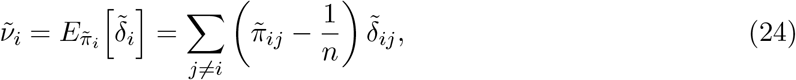

where 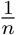 is subtracted to correct for non-uniform density of embedding points. This displacement then represents the embedded velocity vector, which is visualized using a streamplot or a grid [14].

### 4.3 Stream and grid plots

Stream plots of velocities on the embedding are generated using `scVelo.pl.velocity_embedding_stream`. Velocity lengths below a 20-percentile threshold were masked. Given that the number of cells is ~500 for either experiment, a more granular representation for each cell is also represented using `scVelo.pl.velocity_embedding grid` with `n_neighbors` set to 1.

Cluster colors for plots were defined in one of two ways: the implicit `CellType` labels ({Inh, Ex, NeuN+, NeuN−}) and colors using the Louvain community detection algorithm on the nearest neighbor graph (via scVelo). To compare results to a control, the same data was plotted using the deterministic RNA velocity model, calculated using `scVelo.velocity` such that “spliced” and “unspliced” layers were assumed to be 5hmC and 5mC data, respectively.

### 4.4 Feature ranking

Features (represented by 100-Kb bins) were first ranked, and then annotated into respective genes for downstream qualitative analysis, using `scVelo.tl.rank_velocity_genes`. This method uses the Welch t-test on HMCVelo expression, and finds features per cell type showing dynamics that are differentially regulated compared to other cell types. As a validation method, ranked features were explored in the Allen Mouse Brain Atlas [28] for their status as differentiation genes, and Biopython was used to search top Entrez results for papers on each gene.

An implementation of HMCVelo will be available at https://github.com/prmshr/HMCVelo.

## 5 Data

### 5.1 Data sources

Two datasets of Joint-snhmC-seq data from murine cortical neurons (adult mouse brain, 6–8 weeks old) were used for proof-of-concept evaluation. The data was generated at the Wu lab [12]. The most relevant datasets are summarized in Table 1.

**Table 1:** Data inputs for experiments 1 and 2.

| ID | Source | Library strategy | $n$ | Format | Experiment |
| --- | --- | --- | --- | --- | --- |
| A | GSE236784 | Bisulfite-assisted ACE-seq | 565 | .bismark.cov | 1 |
| B | GSE236789 | Bisulfite-Seq | 519 | .bismark.cov | 1 |
| C | In-house | Both (JointSeq) | 545 | .h5ad (Seurat) | 2 |

#### Dataset 1 (data A+B)

519 cells with high-dimensional read counts from bisulfite-assisted ACE-seq (5hmC) and bisulfite-seq (5mC+5hmC) in *>*26,000 100-Kb genomic bins, available under GEO SuperSeries GSE236798. Cells were selected based on mappability between the two sequencing modalities using composite primary keys ⟨*pool_id, barcode_id, seq_date, input_type*⟩, where `input_type` comprised labels {Ex, Inh, NeuN_neg, NeuN_pos}. Data A contained two files per cell (CpH and CpG reads). The full dataset is *>*110 Gb in size, requiring specialized methods for processing.

#### Dataset 2 (data C)

545 cells with normalized proportions for 5hmC and 5mC across 8,649 100-Kb bins, stored as Seurat objects and converted to AnnData format with two corresponding layers (5hmC, 5mC).

Cell type labels comprised: excitatory neurons (Ex), inhibitory neurons (Inh), NeuN-positive (NeuN+, mature), and NeuN-negative (NeuN−, immature).

### 5.2 Preprocessing

#### True mC calculation

scBS-ACE-seq provides subtraction-free and high-resolution counts of 5hmC abundance in nonoverlapping 100-Kb bins across the genome (Fig. 1). While the 5hmC signal was calculated as % of C/(C+T) at each gene through direct reads (for data C) or as the raw read count per bin (for A, B), 5mC was calculated indirectly through subtraction: scBS-seq data − scBS-ACE-seq data.

#### Normalization

Counts-per-cell normalization was conducted for all data using Episcanpy. Highly variable genes were excluded from normalization.

#### Inference of unmodified cytosine for Experiment 2

For data C, 5hmC and 5mC signals are calculated as % of reads/total C reads. JointSeq notes a strong similarity in covered CpG sites per nucleus and uniquely mapped reads per nucleus between both protocols given its matching sequencing depth [12]. Therefore, for experiment 2, it was assumed that relative cytosine proportion *x*_*c*_ for gene *g* and cell *C* can be calculated using the normalized proportions *x*_*m*_ and *x*_*h*_ for 5mC and 5hmC using *x*_*c*_ = 100% − *x*_*m*_ − *x*_*h*_.

#### Gene mapping

UCSC Table Browser [26] was used to convert 100-Kb bins to RefSeq gene IDs using a one-one mapping. Data for features spanning more than one 100-Kb bin were aggregated. Due to the sparsity of methylation data as well as unknowns about the exact location and number of promoters for a gene, binning is often implemented in single-cell methylation studies [27]. A 100-Kb bin is a large enough range such that a gene as well as its promoter’s methylation can be considered.

## 6 Results

Results were computed for the two experiments using two separate datasets. This research was conducted on adult mouse brains (6–8 weeks old). While this makes the steady-state assumption very appropriate, not all cells will have differentiating trajectories, and therefore the results must be taken as a validation of the methodology in resolving a trajectory wherever one exists.

### 6.1 HMCVelo resolves lineage source and branching

Applied to Dataset 1, HMCVelo reveals a bifurcating trajectory originating from NeuN-negative (immature) cells (Fig. 3). In the velocity stream plots (plotted on UMAP embedding of OMS), HMC velocity shows two branches. The source of the directions can be visually inspected as NeuN-negative cells. Using Louvain clusters, the NeuN-negative cluster appears to be resolved into two individual clusters representing transient states moving toward either direction. Additionally, inhibitory and excitatory clusters populate either side of the stream plot, raising the hypothesis that these lineages move toward either inhibitory or excitatory states. NeuN-positive cells populate both branches, consistent with terminal differentiation. Similar branching topology is observed in t-SNE embeddings (Fig. 4).

**Figure 3:**
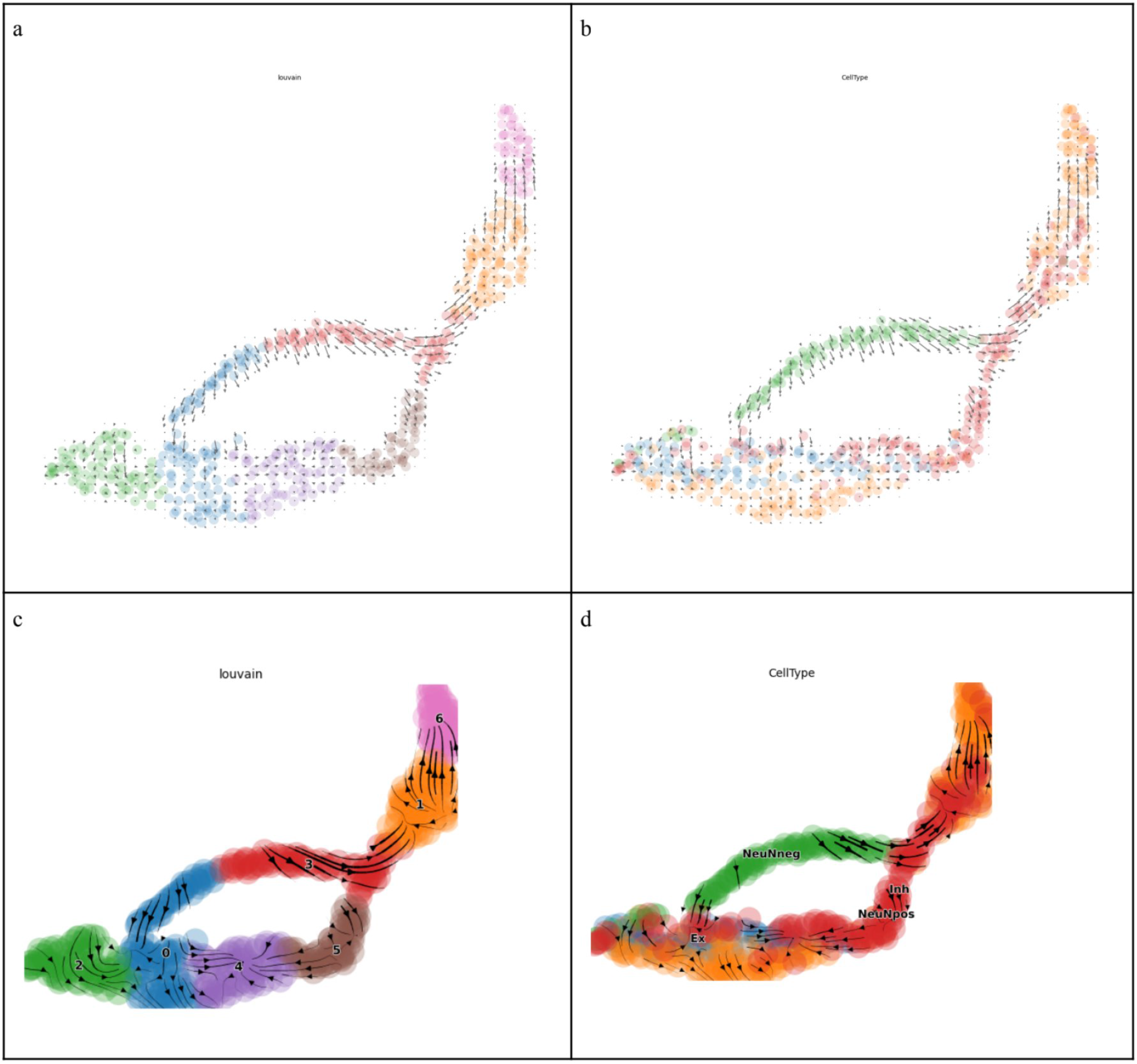
Grid plots (a,b) and stream plots (c,d) using UMAP and Louvain cluster colors for data A,B (a,c); and for CellType clusters (b,d).

**Figure 4:**
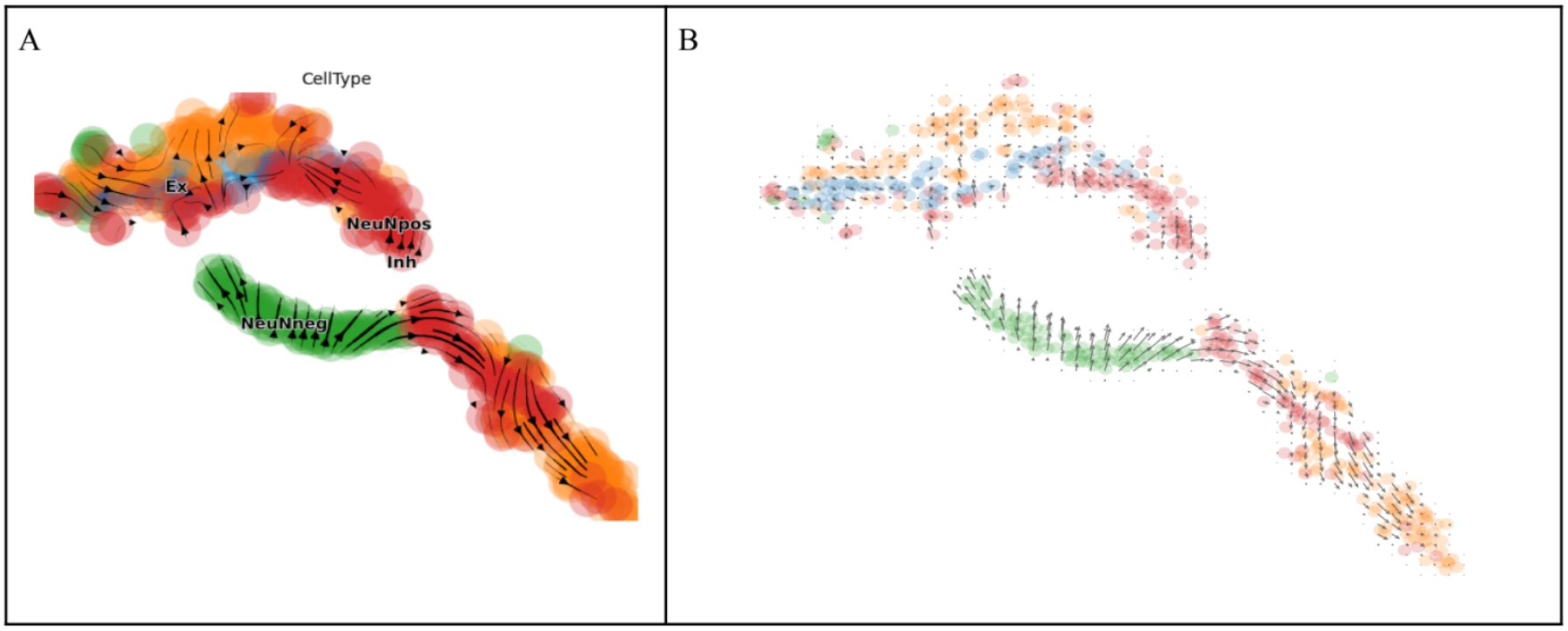
t-SNE also shows divergent directions in stream (a) and grid (b) plots.

**Remark 1** (Choice of embedding basis). Let *x*_*m*_(*t*), *x*_*h*_(*t*), *x*_*c*_(*t*) denote the proportions of 5mC, 5hmC, and unmodified cytosine at gene *g* and cell *i*, subject to the conservation constraint

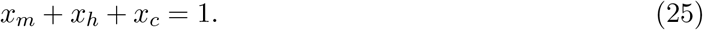

It follows that *x*_*c*_ = 1 − *x*_*m*_ − *x*_*h*_, and thus the unmodified cytosine proportion is not an independent variable but a *derived quantity* —a linear projection of the joint (*x*_*m*_, *x*_*h*_) signal onto a one-dimensional subspace orthogonal to the simplex.

We observe empirically that embedding on *x*_*c*_ collapses the bifurcating trajectory resolved by HMCVelo into a single lineage exhibiting a characteristic horseshoe artifact (Fig. 5). This is not an incidental failure of dimensionality reduction but a *necessary consequence* of the information geometry. The excitatory and inhibitory branches are distinguished by the *covariance structure* between *x*_*m*_ and *x*_*h*_—that is, the branches separate precisely because 5mC and 5hmC redistribute differently across divergent fates. The observed methylation state *x*_OMS_ = *x*_*m*_ + *x*_*h*_ preserves this joint variation. By contrast, *x*_*c*_ = 1 − *x*_OMS_ is a scalar affine transformation of *x*_OMS_ that retains only the total magnitude of modification while discarding the relative allocation between methylation and hydroxymethylation—the very degrees of freedom that encode branching.

**Figure 5:**
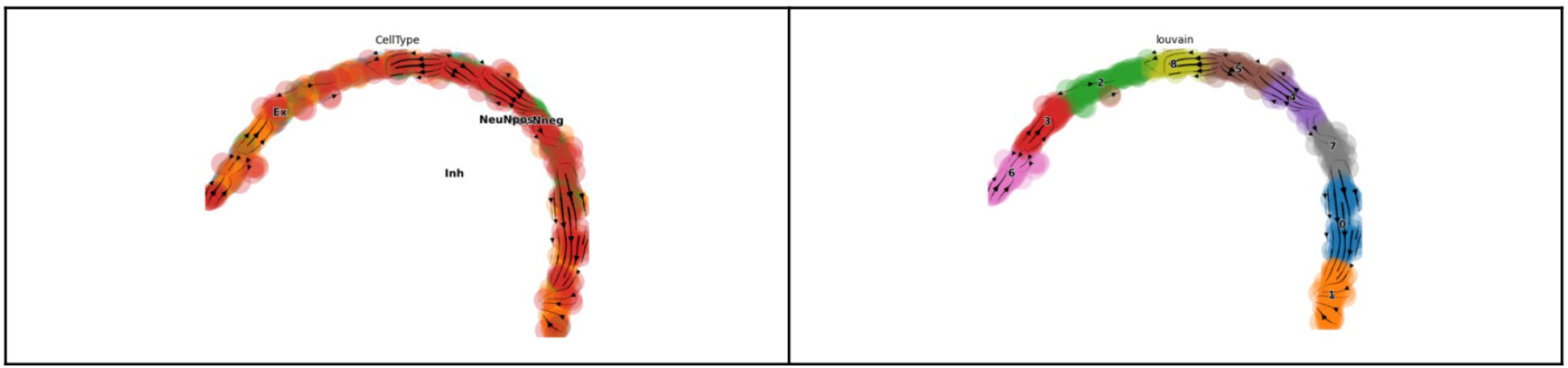
The usage of dimensionality reduction and velocity embedding on unmodified cytosines. Embedding on *x*_*c*_ collapses the bifurcation into a horseshoe artifact, as predicted by Remark 1.

More precisely: let 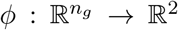 denote a nonlinear embedding (e.g., UMAP). The input vectors 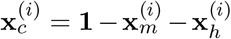 lie in a subspace of dimension rank(**X** + **X**), which is identical to the column space of **X**_OMS_ but reflected. When the intrinsic dimensionality of the biological manifold exceeds the effective dimensionality captured by *x*_*c*_—as occurs at any bifurcation point where the two daughter lineages diverge along axes not aligned with the total modification gradient—the embedding *ϕ*(**x**_*c*_) cannot faithfully represent the fork. The continuous trajectory is projected onto a curve of lower intrinsic dimension, and the well-known horseshoe distortion arises as a generic artifact of such rank-deficient embeddings under nonlinear maps [30, 31].

This result generalizes: in any closed biochemical system with a conservation law ∑_*k*_*x*_*k*_ = const, the complement variable *x*_complement_ = const − ∑_*k≠*=*j*_ *x*_*k*_ is algebraically redundant with the sum of the remaining states and cannot resolve trajectory structure that depends on the *partitioning* among those states. The correct embedding basis for velocity visualization is therefore the joint observed signal—here, *x*_OMS_ = *x*_*m*_ + *x*_*h*_—which retains the full covariance between measured species.

We note that this observation has practical implications beyond the present model. Any future velocity framework applied to cyclic biochemical systems (e.g., histone modification cycles, nucleotide salvage pathways) must similarly attend to the choice of embedding basis, ensuring that the input to dimensionality reduction preserves the independent axes of variation along which cellular trajectories diverge.

### 6.2 HMCVelo outperforms repurposed RNA velocity on methylation data

As a baseline comparison, deterministic RNA velocity (scVelo [14]) was applied to the same data by treating 5hmC as “spliced” and 5mC as “unspliced” counts, plotted over identical embeddings. HMCVelo achieves substantially higher velocity confidence across all cell types (Table 2).

**Table 2:**
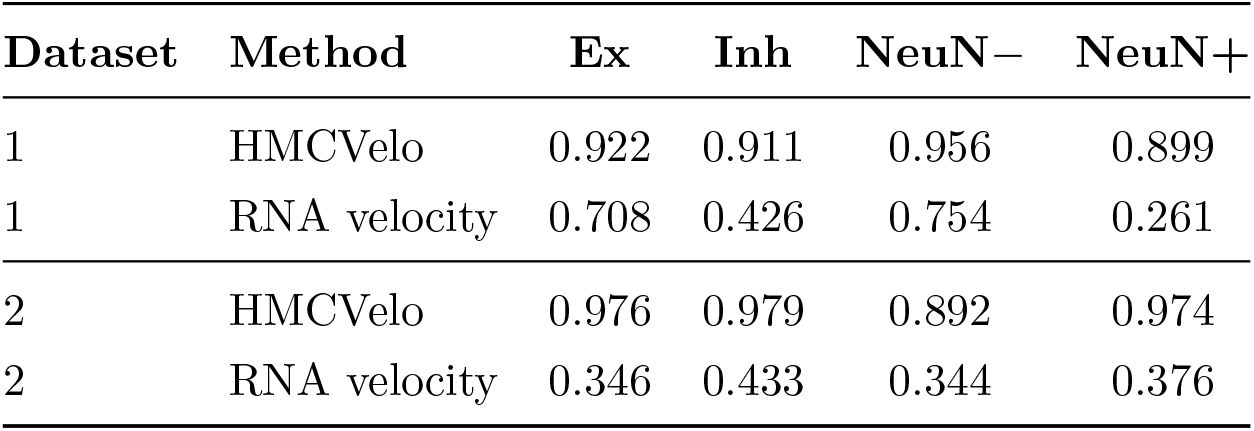
Velocity confidence by cell type for HMCVelo versus repurposed RNA velocity.

On Dataset 1, mean velocity confidence for HMCVelo ranges from 0.90 to 0.96 across cell types, compared to 0.26 to 0.75 for RNA velocity. On Dataset 2, HMCVelo achieves confidence scores of 0.89 to 0.98, compared to 0.34 to 0.43 for RNA velocity. Qualitatively, RNA velocity grid plots show disorganized, locally incoherent arrows (Fig. 6a), whereas HMCVelo produces smooth, globally consistent flow fields (Fig. 6b).

**Figure 6:**
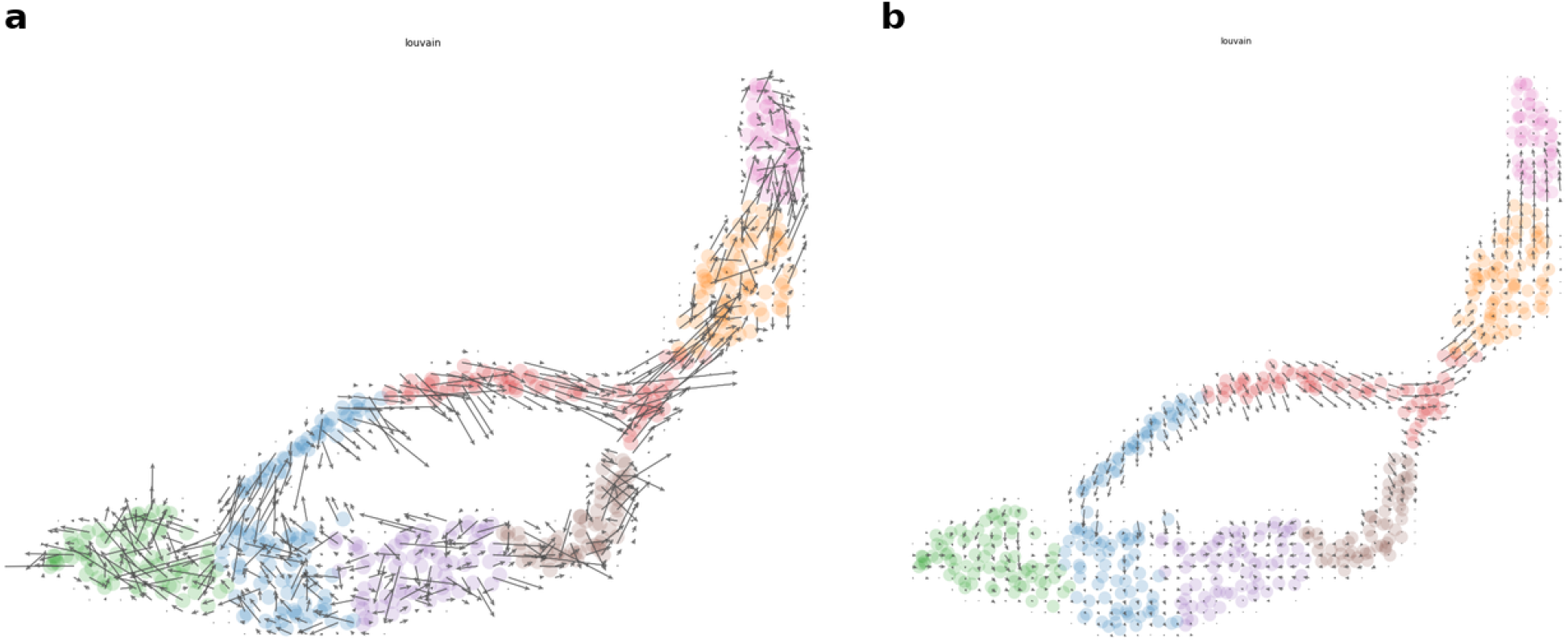
Velocity grid plots on identical UMAP embeddings of observed methylation states. (a) Repurposed RNA velocity (deterministic mode, scVelo). (b) HMCVelo. Arrows in (a) are locally incoherent, while (b) shows smooth, globally consistent flow fields with clear bifurcation from immature to excitatory and inhibitory lineages.

**Figure 7:**
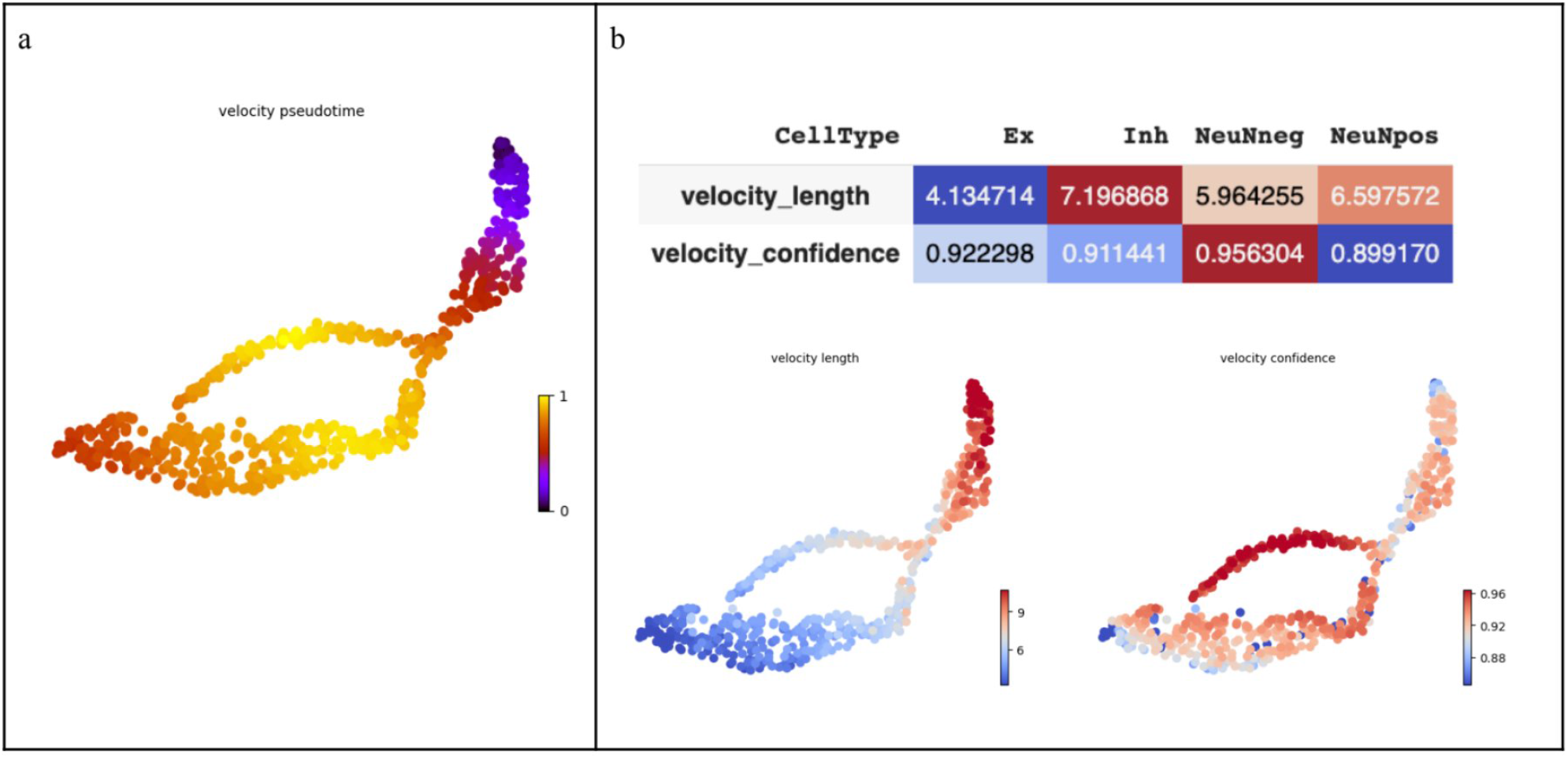
HMCVelo pseudotime and velocity confidence on Dataset 1. (a) Velocity pseudotime shows a smooth gradient from NeuN− (root, purple) to terminal states (yellow). (b) Velocity confidence exceeds 0.89 across all cell types, with velocity length and confidence spatial maps shown below.

**Figure 8:**
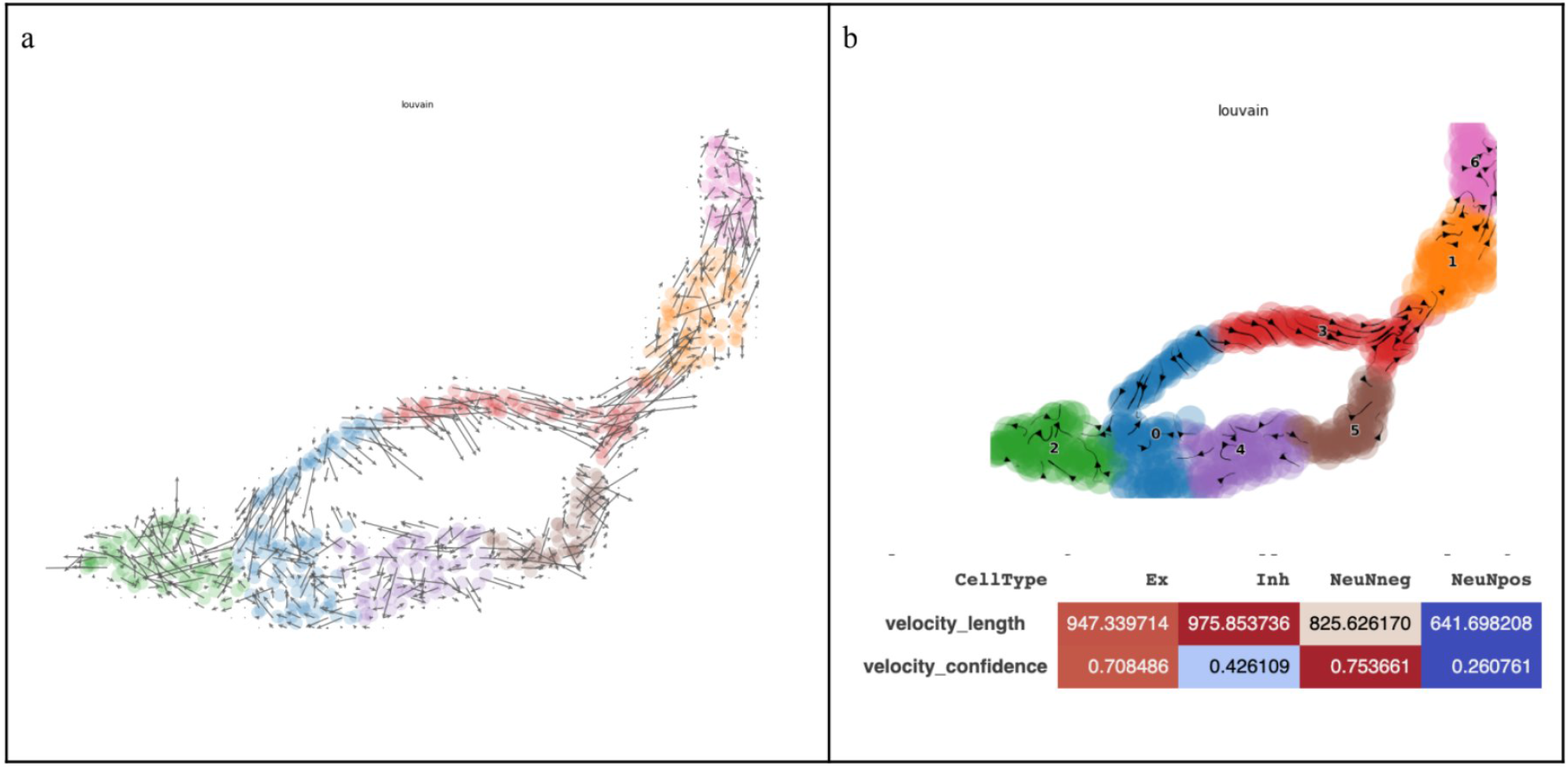
Repurposed RNA velocity on identical data (Dataset 1). (a) Grid plot shows disorganized, locally incoherent arrows. (b) Stream plot with Louvain clusters and velocity confidence table. Velocity confidence drops to 0.26–0.75, with NeuN+ cells at 0.26. Note the order-of-magnitude difference in velocity length scales compared to HMCVelo (Fig. 7).

### 6.3 Ranked velocity genes implicate known differentiation regulators

Genes were ranked by differential velocity dynamics across cell types using the Welch t-test (via `rank_velocity_genes` in scVelo, minimum correlation threshold 0.3). For RNA velocity, this procedure returned no genes meeting the threshold—further evidence that the repurposed model fails to capture meaningful dynamics in methylation data.

For HMCVelo, the top-ranked genes from Experiment 1 include:

> *Gprc5d, Cyp2c38, Ddx47, Fam234b, Cd200*, ***Pde1c, Afap1, Lingo1, Cry2, Grid2, Sntg2, Mapk8ip1, Ski, Satb2, Atf7ip***, *Morn1, Ccser1, Myh7b, Sfi1*, …

Genes in bold were independently corroborated through the Allen Mouse Brain Atlas [28] and literature search. *Satb2* is a transcription factor critical for callosal projection neuron identity and cortical layer specification. *Lingo1* regulates myelination and neurite outgrowth. *Grid2* encodes an ionotropic glutamate receptor involved in synaptic development. *Mapk8ip1* (JIP1) scaffolds JNK signaling in axonal transport. *Cry2* functions in circadian regulation with emerging roles in neural development.

Top-ranked genes from Experiment 2 include:

> *Dynll1, Tacc1, Kalrn, Scn8a, Dlgap4, Nedd4l, Bsn, Slc6a7*, ***Prkcb, Camk2a, Stmn1, Satb2***, *Calm1, Sptbn4, Nav2, Gas7, Prkce*, …

*Camk2a* and *Prkcb* are kinases central to synaptic plasticity. *Stmn1* is involved in microtubule dynamics during neuronal differentiation. *Satb2* appears in both experiments, supporting the robustness of the velocity ranking. See Supplementary Data for complete gene lists and literature validation.

**Figure 9:**
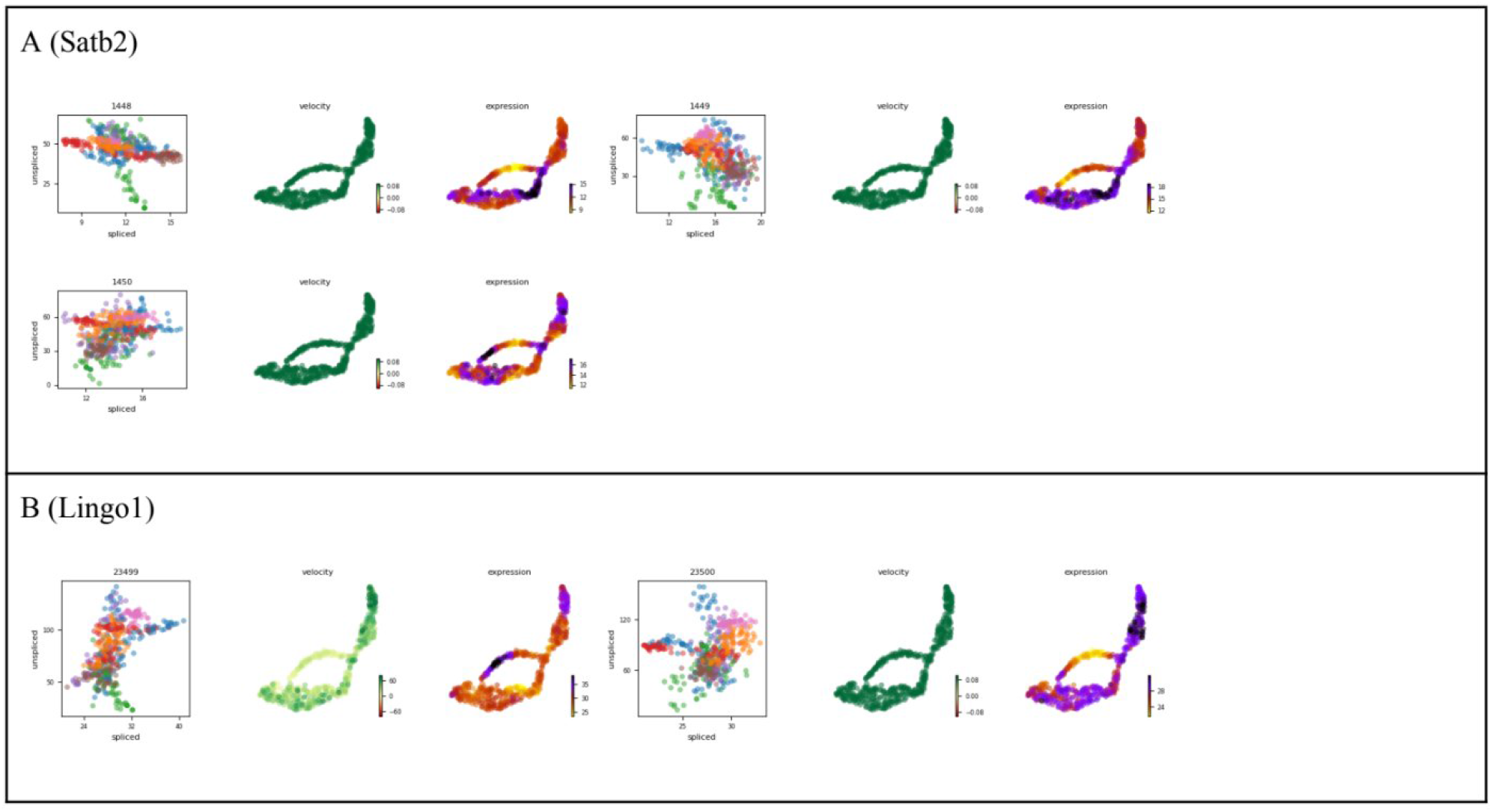
Velocity plots for top-ranked genes (part 1 of 3). Each gene shows: phase portrait (5hmC vs. 5mC, colored by Louvain cluster), velocity magnitude on embedding, and expression on embedding. (A) *Satb2* : callosal projection neuron identity and cortical layer specification. (B) *Lingo1* : myelination and neurite outgrowth. “Spliced” refers to 5hmC; “unspliced” to 5mC.

**Figure 10:**
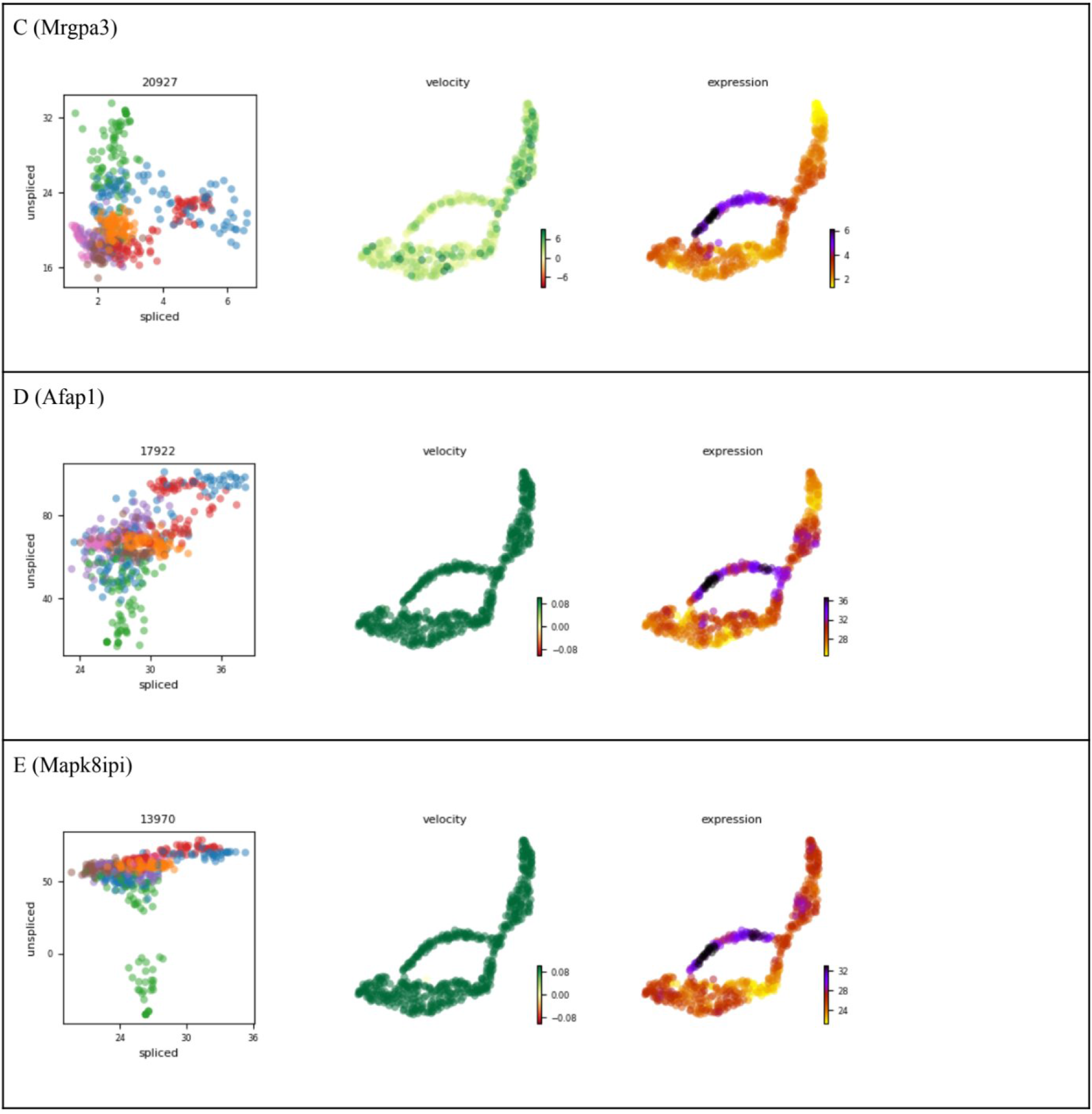
Velocity plots for top-ranked genes (part 2 of 3). (C) *Mrgpra3* : sensory neuron marker. (D) *Afap1* : actin filament-associated protein. (E) *Mapk8ip1* (JIP1): JNK signaling scaffold in axonal transport.

**Figure 11:**
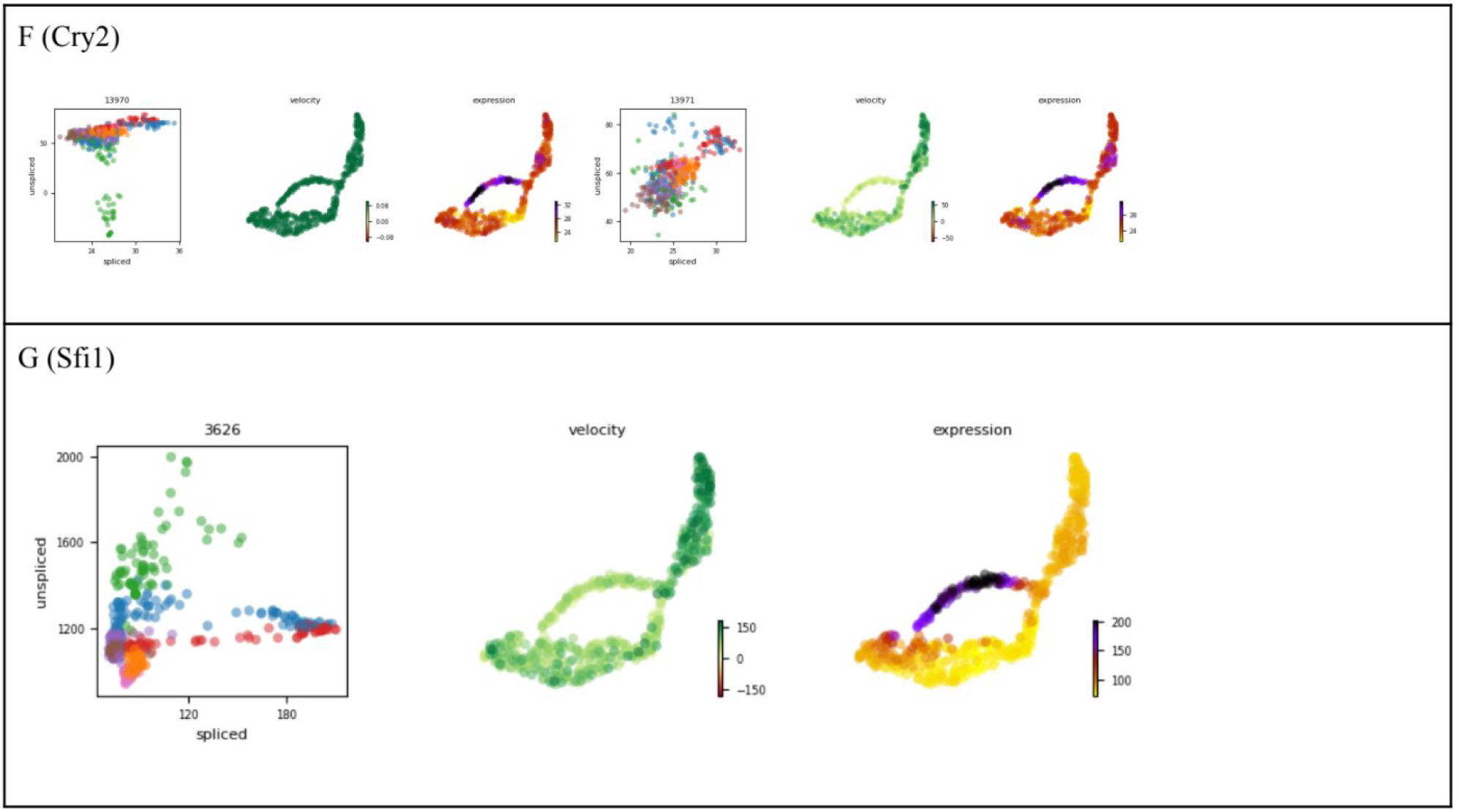
Velocity plots for top-ranked genes (part 3 of 3). (F) *Cry2* : circadian regulation with emerging roles in neural development. (G) *Sfi1* : centrosome dynamics during cell division.

### 6.4 Experiment 2: Hydroxymethylation velocity on mature and immature cells

While there is significant whitespace and difference in cell clusters between mature and immature cells for Dataset 2, there is clear directionality in the data. From the pseudotime calculations (Fig. 13), HMCVelo successfully reveals the root (NeuN−) and terminal clusters. While this data is more processed and has fewer genes than Dataset 1, it serves as a relevant proof-of-concept for HMCVelo on proportion-based data.

**Figure 12:**
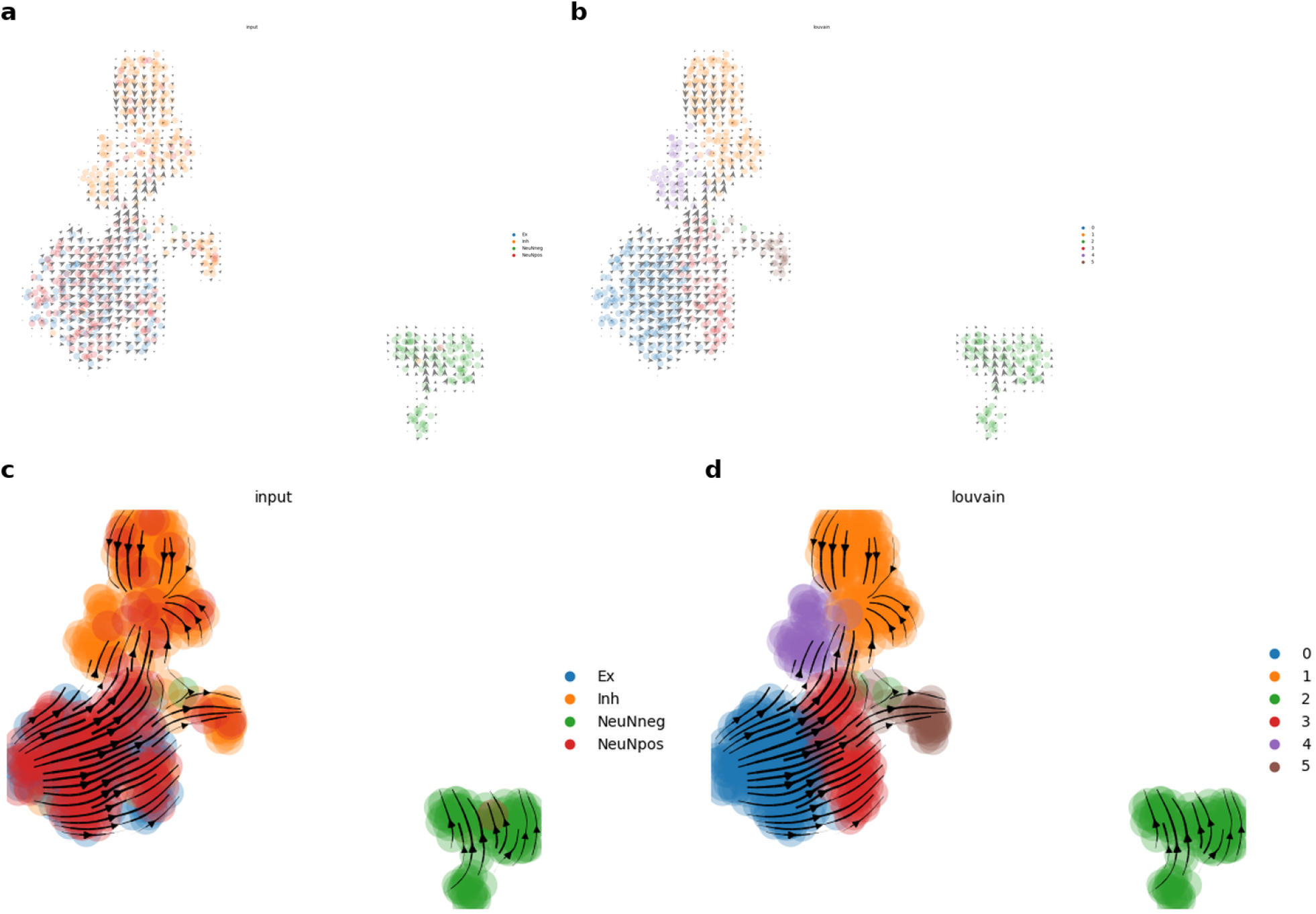
Experiment 2: HMCVelo applied to Dataset 2 (545 cells, proportion-based data). Grid plots (a,b) and stream plots (c,d) using UMAP for CellType clusters (a,c) and Louvain clusters (b,d). Clear directionality from NeuN− (immature) toward mature cell states is visible despite the whitespace between clusters.

**Figure 13:**
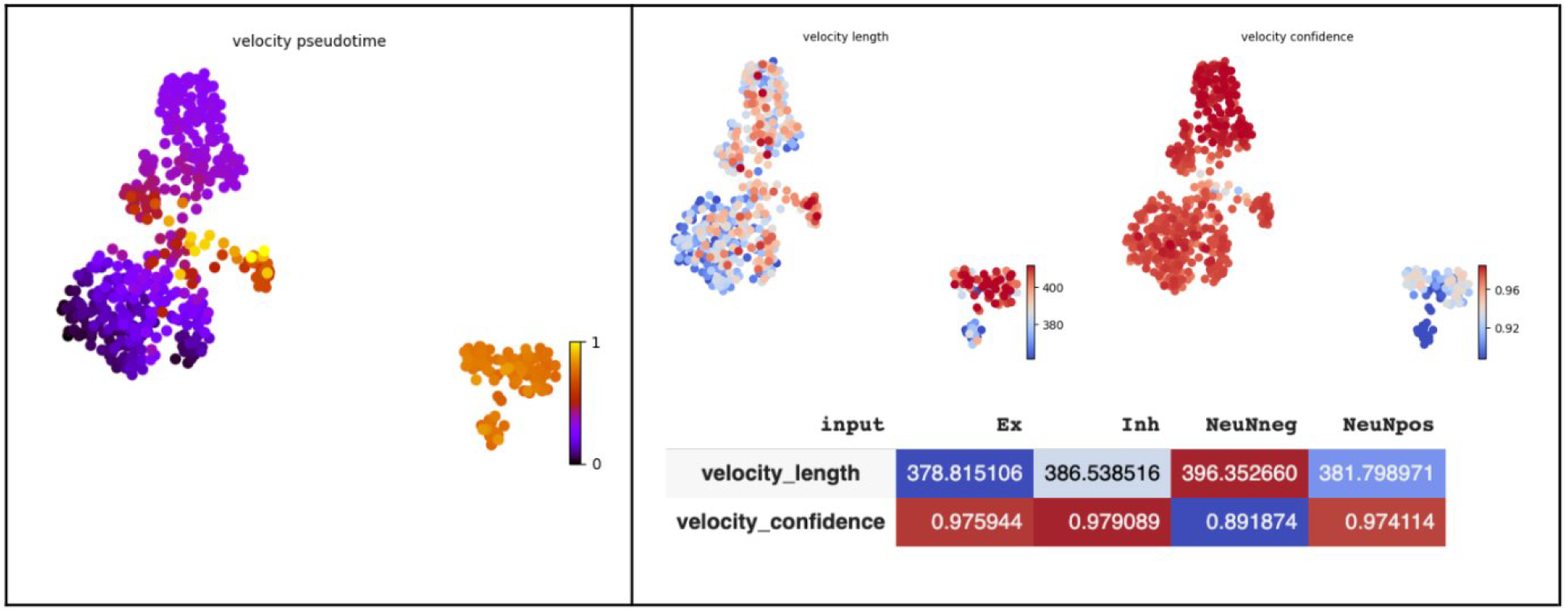
Experiment 2: HMCVelo pseudotime and velocity confidence. (a) Velocity pseudotime correctly identifies NeuN− as root (purple) with progression toward terminal states. (b) Velocity confidence exceeds 0.89 across all cell types (yellow), confirming robust trajectory inference on proportion-based data.

## 7 Discussion

This is the first attempt at calculating a methylation-based velocity vector. By modeling the methylation–demethylation cycle as a system of coupled ODEs with gene-specific rate parameters, the method extracts directional information from static molecular snapshots that is inaccessible to expression-based models repurposed for epigenetic data. The substantial improvement in velocity confidence over RNA velocity applied to the same data (Table 2) demonstrates that a model grounded in the biochemistry of the methylation cycle—specifically, the cyclic rather than unidirectional nature of the process and the inclusion of unmodified cytosine—captures dynamics that a two-variable linear model cannot.

### Limitations and future directions

There are several limitations to the model that define clear directions for extension.

First, HMCVelo relies on qualitative validation, as Joint-snhmC-seq is a recently developed protocol and no comparable 5hmC/5mC single-cell datasets exist for benchmarking against established trajectory inference methods. In the medium-term, through the generation of Joint-snhmC-seq data alongside mapped mRNA reads from scNuc-drop-seq, it may be possible to directly compare the theoretical HMC-RNA velocity discussed above to traditional RNA velocity on the same data. While it has been deduced in the past that 5hmC values partially predict transcription, this could provide a true single-cell estimation of the prediction.

Second, RNA velocity uses distinguishable features to resolve unspliced vs. spliced counts (e.g., reads spanning exon/intron vs. exon/exon boundaries). Meanwhile, JointSeq data relies on two sequencing strategies with different sensitivities; while data from samples can be mapped to assume parallel datasets, this is a noisy assumption, and more protocol-specific preprocessing techniques could be applied to reduce noise.

Third, the steady-state assumption may fail for transiently expressed genes or for populations lacking terminal cell states. A dynamical extension of the model, analogous to scVelo’s dynamical mode [14], could relax this assumption. Crucially, RNA velocity uses an analytical solution for ODEs, while HMCVelo uses a numerical solution relying on the time-step chosen for analysis.

Fourth, 100-Kb bins may be too coarse to resolve promoterversus gene body–specific methylation dynamics. By distinguishing promoter and non-promoter methylation, more insights can be obtained; this is especially possible since JointSeq already distinguishes CpH and CpG methylation.

Fifth, passive demethylation is not captured and would require additional data, potentially from cell-cycle-resolved experiments. It may be possible to make more inferences about passive methylation using feature engineering on genes relevant to DNA methylation maintenance, most suitably using gene expression data.

Sixth, it may be overly simplistic to describe hydroxymethylation dynamics with one universal parameter, given there are noted instances where a cytosine remains hydroxymethylated versus completing the demethylation cycle. A switch parameter could be used in a future implementation, but the switch time for such a parameter requires more foundational knowledge on 5hmC in the brain.

Finally, similar to RNA velocity, given the high-dimensionality of the velocity vector, the final plotting is highly dependent on the underlying embedding algorithm, the number of features chosen, and the layer used for embedding. Remark 1 formalizes one aspect of this sensitivity. There can be multiple non-dependent components for various processes like maturation and differentiation, which are masked in low-dimensional representations. Newer approaches to tailor embeddings can be considered.

Despite these limitations, HMCVelo provides a principled framework for extracting temporal information from the epigenome. As joint methylation–transcription profiling technologies mature, this approach can serve as a component of multi-omic velocity models that integrate dynamics across the central dogma—from DNA methylation to transcription to protein—enabling a more complete characterization of the forces governing cellular differentiation.

## Acknowledgments

I thank Dr. Hao Wu and the Wu laboratory (Department of Genetics, Perelman School of Medicine and Penn Epigenetics Institute, University of Pennsylvania) for providing the Joint-snhmC-seq data [12] used in this study and for guidance on single-cell epigenomics. I thank Dr. Joshua B. Plotkin (Department of Biology, University of Pennsylvania) for overseeing the independent study under which this work was initiated.

## Data Availability

Raw sequencing data for Dataset 1 are available under GEO SuperSeries GSE236798. Dataset 2 was generated in-house at the Wu lab. An implementation of HMCVelo will be available at https://github.com/prmshr/HMCVelo. Annotated Colab notebooks reproducing all analyses are available on request.

## Supplementary Figure

**Figure 14: Supplementary Figure 1.**
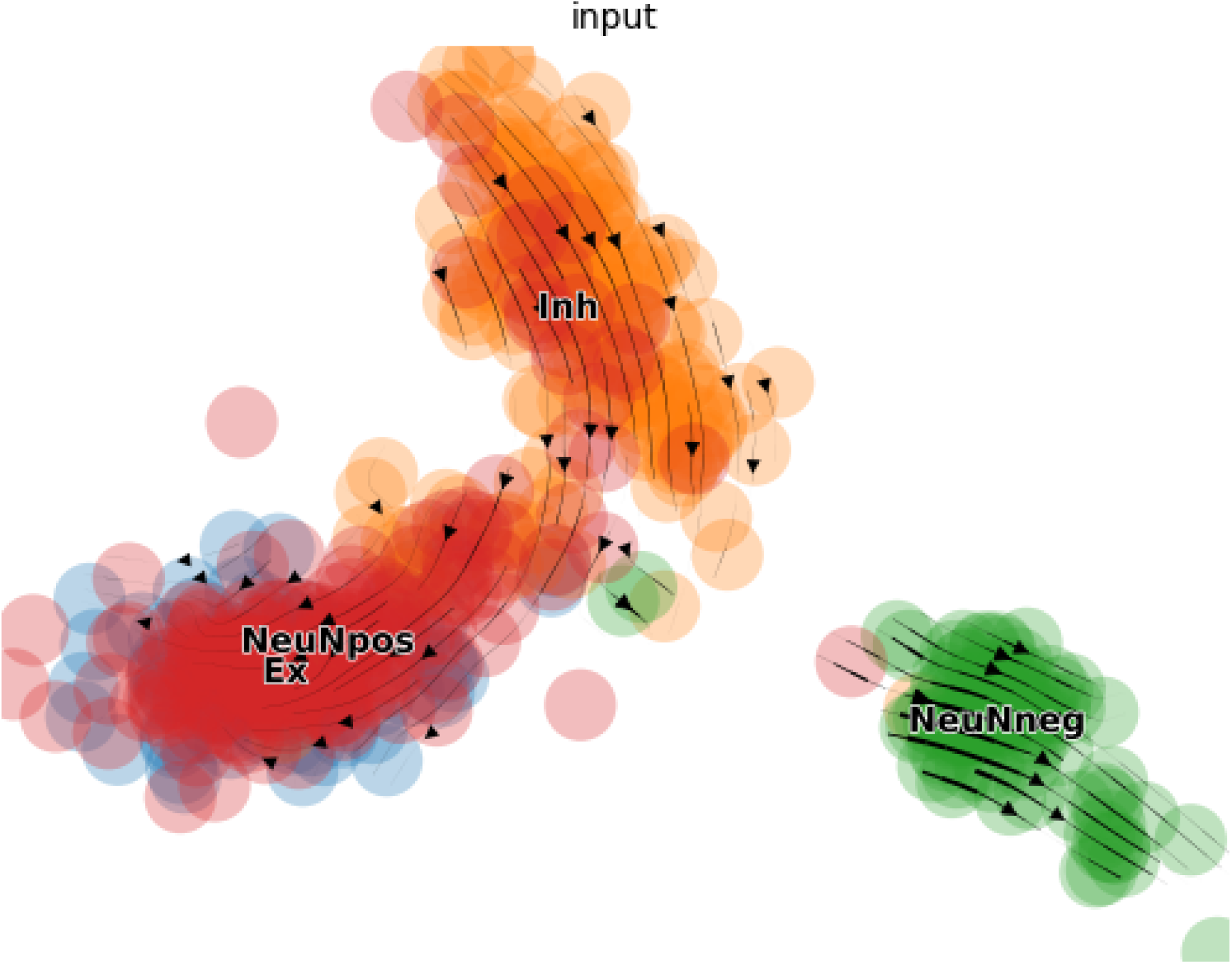
HMCVelo velocity stream on PCA embedding for Dataset 2, colored by cell type. The PCA basis resolves the three clusters (Inh, NeuNpos/Ex, NeuNeg) with clear within-cluster directionality, confirming that HMCVelo trajectory inference is robust across embedding methods.

